# WBSCR16 Determines Metabolic Flexibility Via Regulating Mitochondrial 16S rRNA Maturation

**DOI:** 10.1101/2023.06.15.545185

**Authors:** Shengjie Zhang, Zi Dong, Yang Feng, Wei Guo, Chen Zhang, Yifan Shi, Zhiyun Zhao, Jiqiu Wang, Guang Ning, Guorui Huang

**Affiliations:** Department of Endocrine and Metabolic Diseases, Shanghai Institute of Endocrine and Metabolic Diseases, Ruijin Hospital, Shanghai Jiaotong University School of Medicine, Shanghai, 200025, China; Shanghai National Clinical Research Center for Metabolic Diseases, Key Laboratory for Endocrine and Metabolic Diseases of the National Health Commission of the PR China, Shanghai Key Laboratory for Endocrine Tumor, Ruijin Hospital, Shanghai Jiaotong University School of Medicine, Shanghai, 200025, China; National Research Center for Translational Medicine, State Key Laboratory of Medical Genomics, Ruijin Hospital, Shanghai Jiaotong University School of Medicine, Shanghai, 200025, China

## Abstract

Mitochondria play significant roles in energy homeostasis by dissipating excess calories. However, the key proteins in mitochondrial govern metabolically flexibility remain poorly understand. Herein, we generated adipose specific *Wbscr16^-/-^*mice and cells, both of which exhibited a preference for fat utilization and displayed dramatic mitochondrial changes. We demonstrated that WBSCR16 is essential for the specific cleavage of 16S rRNA-tRNA^Leu^, leading to the production of maturated 16S rRNA, which in turn controls mitochondrial large ribosomal assembly. Importantly, WBSCR16 was identified as a 16S rRNA-binding protein that directly interacts with RNase P subunit MRPP3. Furthermore, evidence showed that WBSCR16 ablation promotes energy wasting via lipid cycling in BAT, leading to excess energy expenditure and resistance to obesity. Conversely, cells and mice with WBSCR16 overexpression preferred to glucose utilization. These findings suggest that WBSCR16 is the key mitochondrial switch protein between glucose and lipid metabolism, making it a promising target for the development of fat busting drug.

## Introduction

More than 70% of the Americans and 50% of the Chinese individuals suffer from overweight and obesity, and are expected to increase in the coming decades ^1^. Adipose tissue plays a crucial role in energy regulation through fat storage and inter-organ communication. Brown adipose tissue (BAT), characterized by its high vascularity organ with abundant mitochondria, dissipates chemical energy stored in lipid droplets through fatty acid (FA) oxidation or thermogenesis, thereby regulate whole-body energy homeostasis ^1, 2^. In addition to utilizing triglyceride stores, BAT oxidizes circulating free fatty acid (FFA) and glucose taken up from the plasma to generate heat for optimal thermogenesis ^3, 4^. BAT produces heat through non-shivering thermogenesis. As a result, therapeutic approaches targeting BAT have gained interest as potential treatments for obesity in humans^5, 6^. Metabolic flexibility, the ability of cells to switch between glucose and lipid fuels, was observed under different nutrient availability conditions and is often accompanied by insulin resistance and mitochondrial dysfunction ^7, 8^. Modulating mitochondrial energy expenditure in adipose tissues is critical for regulating energy homeostasis, insulin sensitivity, glucose and fat metabolism, and thermogenesis ^9–11^. But little is known about which mitochondrial proteins are involved and how they manipulate metabolic flexibility and substrate preference.

Mitochondria is a vital organelle for metabolic activities, produce the majority of ATP through oxidative phosphorylation (OXPHOS) via mitochondrial electron transport chains (ETCs) ^12, 13^. In addition to being regulated by nuclear-encoded protein factors, ETC functions are governed by the expression of mitochondrial genes ^14–16^. Mitochondrial RNA processing involves cleavage at the 5’ end of tRNAs by RNase P and cleavage at 3’ end of tRNAs by RNase Z ^17, 18^. RNase P is a special mitochondrion-targeted RNase composed of three protein subunits without an RNA component: mitochondrial RNase P protein 1 (MRPP1), MRPP2, and MRPP3 ^17, 19^. MRPP1, MRPP3 and ELAC2 all contain RNA-binding protein domains (RBDs), and current understanding suggests their direct involvement in regulating mitochondrial RNA processing ^18, 20, 21^. While proteins involved in the specific maturation of mitochondrial 12S rRNA, such as TFB1M and NSUN4, have been identified ^22, 23^, those specifically involved in 16S rRNA processing are poor understood.

In our previous study, we identified WBSCR16 as a mitochondrial protein encoded by the nuclear gene *Rcc1L*, functioning as a guanine nucleotide exchange factor (GEF) for the mitochondrial fusion GTPase OPA1^24^. Recent studies have also revealed its roles in regulating mitochondrial gene expression, possibly due to its involvement in mitochondrial ribosome assembly ^25–27^. However, the precise mechanism by WBSCR16 involves in mitoribosomal assembly and its physiology function yet unknown.

In this study, we found that WBSCR16 selectively binds to unprocessed transcripts containing 16S rRNAs through its RBD and interacts with MRPP3 to facilitate the enzyme activity of RNase P, leading to cleavage at the 5’ end of tRNA^Leu^, and subsequent produce mature 16S rRNA. Given the abundance of mitochondrial and energy consumption capacity of BAT, the genetic ablation of WBSCR16 in adipose tissues promotes a metabolic substrates preference from glucose to FA, ultimately enhancing whole-body energy expenditure and conferring resistance to diet-induced obesity. Therefore, targeting WBSCR16 could potentially serve as a novel strategy to increase energy expenditure through efficient lipid cycling, meanwhile avoiding hypoglycemia or hyperglycemia events, offering potential benefits in combating obesity and metabolic diseases.

## Results

### The Alterations of Mitochondrial in *WBSCR16^-/-^* Tissues and Cells

Whole-body deficiency of WBSCR16 causes early embryonic lethality and mitochondrial fragmentation^28^. BAT, for instance, derives its color from the abundance of mitochondria within its cells^29^. The high abundance of mitochondria in BAT regulate energy expenditure by substrate oxidation and thermogenesis, a potential therapeutic target for obesity and diabetes^30^. To investigate the mechanism underlying mitochondrial alterations in adipose and to determine the physiology function of mitochondrial protein WBSCR16 *in vivo*, adipocyte-specific WBSCR16 knockout (AWBSCR16 KO) mice with Adiponectin^Cre^ were generated. As expected, WBSCR16 expression was greatly reduced in AWBSCR16 KO iBAT and WAT, but remain unchanged in the liver (Fig. 1A). Interestingly, an increase in relative iBAT weight and larger lipid droplet size were found in AWBSCR16 KO mice compared to Floxed controls (Fig. 1B-D). Additionally, a decreased in WAT weight was observed, result in a lower fat/lean ratio (Fig. 1E-G). But no differences were observed in overall body weight, WAT lipid size, liver, plasma insulin levels, and basal metabolic parameters such as oxygen consumption in mice between AWBSCR16 KO mice and controls (Fig. S1).

**Fig. 1.**
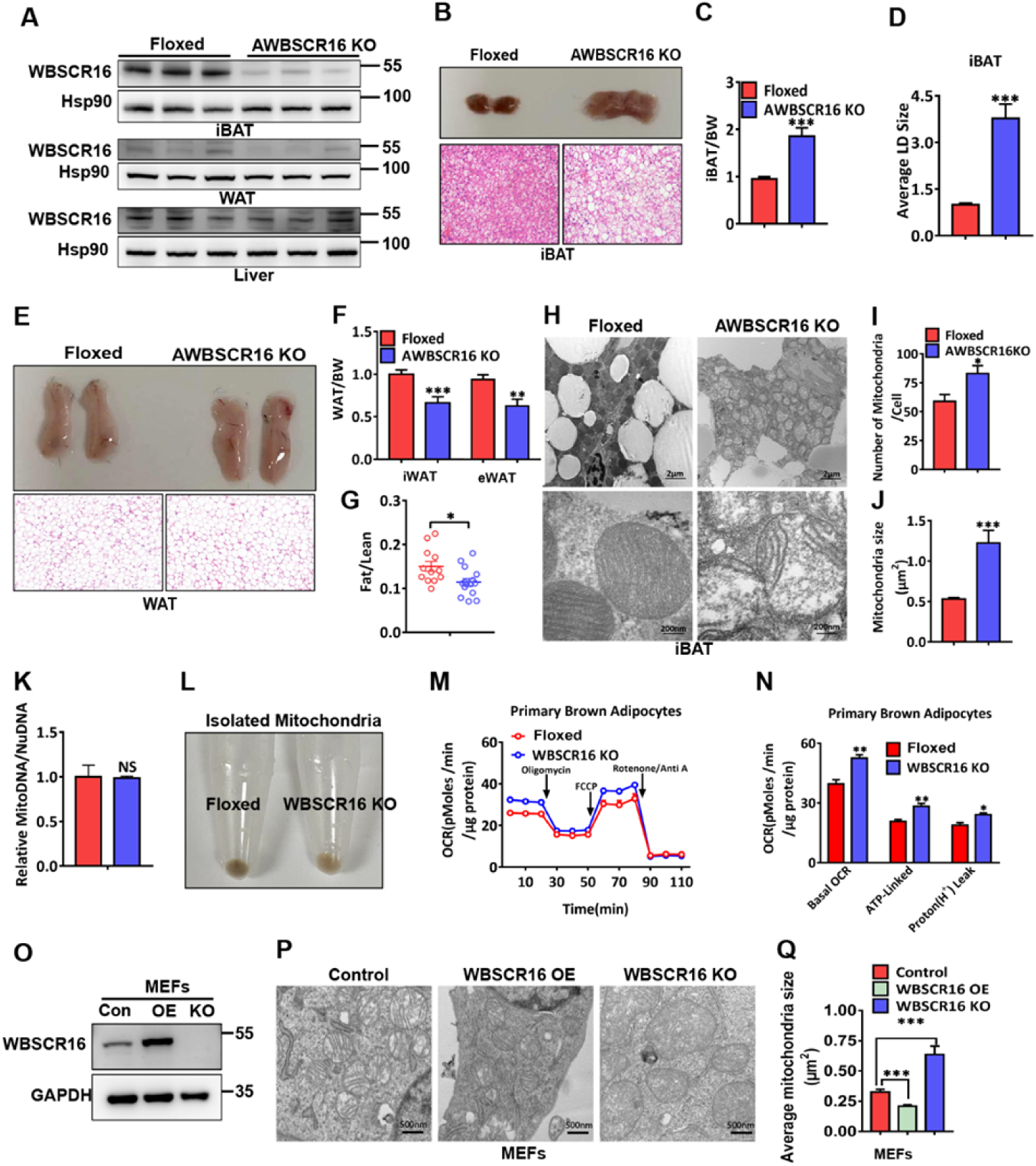
Mitochondrial changes in WBSCR16 null cells and adipose tissues. **(A)** Representative Western blots of WBSCR16 in iBAT, WAT and liver from Floxed and AWBSCR16 KO mice. **(B-D)** Representative pictures and HE staining of iBAT **(B)**, relative weight normalized to body weight **(C)** and cross-sectional area of lipid droplets in iBAT **(D)**. **(E-G)** Representative pictures and HE staining of WAT **(E)**, relative weight normalized to body weight **(F),** and fat mass to lean mass ratio **(G)**. **(H-K)** Electron microscopy images of iBAT **(H)**, number **(I)**, size **(J)** of mitochondria, and the relative mitochondrial DNA content **(K)** in iBAT from Floxed and AWBSCR16 KO mice. **(L)** Representative pictures of mitochondria extracted from iBAT. **(M-N)** Oxygen consumption rate (OCR) of oligomycin, FCCP and antimycin A/Rotenone-treated matured adipocytes derived from the iBAT-SVF of Floxed mice infected with or without Cre lentivirus (**M**), and the quantification of OCR at different stages (**N**). **(O)** Representative western blots of WBSCR16 in overexpression (OE) or knockout (KO) mouse fibroblasts. **(P-Q)** Electron microscopy images (**P**) of MEFs, size (**Q**) of mitochondria in WBSCR16-OE and KO cells.

In comparison to Floxed controls, AWBSCR16 KO iBAT showed dramatically changes in mitochondrial morphology including a substantial reduction in tubular cristae, increased number, and larger size of mitochondria (Fig. 1H-J). However, the copy number of mitochondrial DNA (mtDNA) remained similar between the groups (Fig. 1K). The mitochondria contain a significant amount of iron, which imparts the brown coloration to the fat ^29^. Data showed a lighter grey appearance of mitochondrial crude extract in AWBSCR16 KO mice compared to the darker brown color in controls (Fig. 1L). These remarkable mitochondrial changes in WBSCR16 ablation indicate a critical role for WBSCR16 in mitochondrial maintenance. Surprisingly, primary BAT cells derived from AWBSCR16 KO mice exhibited higher oxygen consumption rates as measured by mitochondrial stress assays using a Seahorse analyzer. This increase was observed at all three stages, including basal respiration, ATP production, and proton leak (Fig. 1M-N). These findings suggest that the decrease in cristae structure in AWBSCR16 KO adipocytes is compensated by an increase in the number and size of mitochondria. Overall, our results demonstrate significant alterations in mitochondrial morphology and function upon WBSCR16 knockout, indicating the crucial role of WBSCR16 in maintaining mitochondrial integrity and regulating energy metabolism in adipose tissues.

To confirm that the changes observed in adipocytes were directly caused by the loss of WBSCR16 and not secondary effects *in vivo*, we established WBSCR16 knockout or overexpression in mouse embryonic fibroblast (MEF) cells (Fig. 1O). Consistently, the mitochondrial morphology in MEF cells was dramatically altered, including changes in mitochondrial size and the number of tubular cristae, depending on the expression of WBSCR16 (Fig. 1P-Q). Surprisingly, despite the loss of mitochondrial cristae, WBSCR16 knockout cells exhibited better mitochondrial membrane potential (Fig. S1K), which is consistent with the higher oxygen consumption observed in AWBSCR16 KO adipocytes (Fig. 1M). This suggests that the defects in mitochondrial morphology were compensated for by an increase in the number, size and membrane potential of mitochondria in WBSCR16-deficient cells. These additional analyses provide further insights into the relationship between AWBSCR16 knockout, mitochondrial morphology, and functional changes in adipocytes and MEF cells.

### Mitoribosomal Assembly Defects in *Wbscr16^-/-^* Cells

Since WBSCR16 deficiency in adipose tissues or MEFs led to a dramatic loss of mitochondrial cristae, we detected certain proteins in respiratory chain complexes and observed a significant decrease in most proteins encoded by the mitochondrial genome (Fig. 2A). Simultaneously, other proteins encoded by the nuclear genome, such as NDUFA4 and NDUFB8 in complex I, and COX IV in complex IV, were also reduced (Fig. 2B). Furthermore, mRNA-coded proteins, two rRNAs, 12S and 16S rRNA, are also encoded by the mitochondrial genome. The amounts of 16S rRNA decreased significantly in AWBSCR16 KO iBAT, while 12S rRNA showed a slight decrease at the same time (Fig. 2C). However, there were no decreases observed in the mRNA levels coding for the above proteins in AWBSCR16 KO iBAT (Fig. 2D). In addition to Since 16S rRNA plays a key role in mitochondrial ribosome assembly, along with other ribosomal proteins, the mitochondrial ribosomal proteins were then investigated. We found an increase in the small ribosomal subunit components MRPS35 and MRPS16 in AWBSCR16 KO iBAT, but no significant change in the large ribosomal subunit components MRPL37 and MRPL12 (Fig. 2E).

**Fig. 2.**
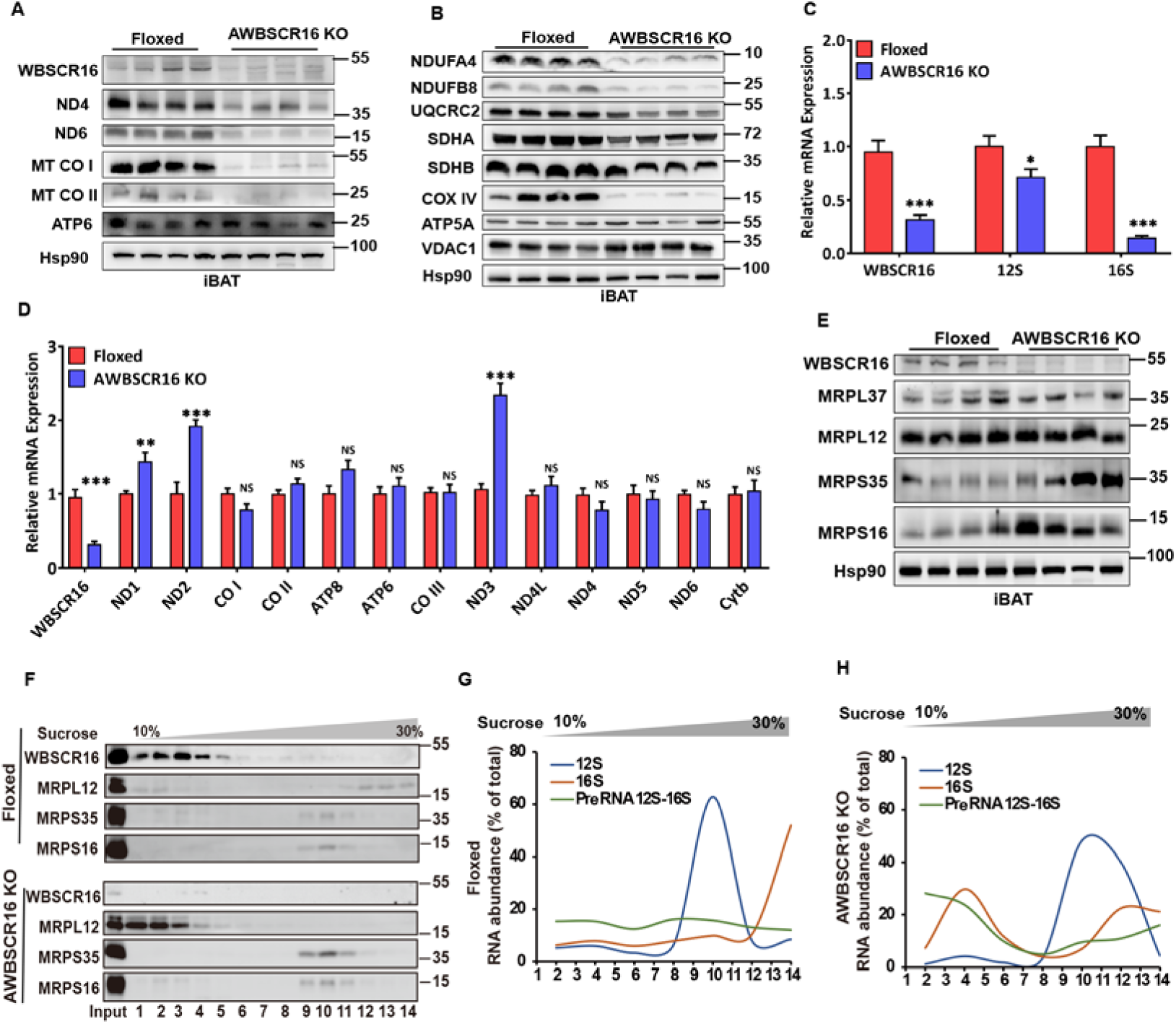
Mitochondrial ribosomal assembly defects in *Wbscr16^-/-^* adipose. **(A)** Representative Western blots of mtDNA encoded protein ND4, ND6, MT COI, MT COII and ATP6, as well as nuclear DNA encoded mitochondrial proteins NDUFA4, NDUFB8, UQCRC2, SDHA, SDHB, COX IV, ATP5A and VDAC1 (**B**) in iBAT. **(C)** qRT-PCR analysis of WBSCR16 and mtDNA encoded RNAs in iBAT from Floxed and AWBSCR16 KO mice. **(D)** qRT-PCR analysis of WBSCR16 and 12S,16S RNAs in iBAT from Floxed and AWBSCR16 KO mice. **(E)** Representative Western blots of WBSCR16, the large ribosomal subunits MRPL37 and MRPL12, and the small ribosomal subunits MRPS35 and MRPS16 in iBAT from Floxed and AWBSCR16 KO mice. **(F)** The distribution of the small and large mitochondrial ribosomal subunits and the monosome in iBAT assayed with continuous 10%–30% sucrose gradients. WBSCR16 and the large ribosomal subunits MRPL37 and MRPL12, as well as the small ribosomal subunits MRPS35 and MRPS16, were detected by immunoblotting with specific antibodies. **(G-H)** Analysis of the distribution of mitochondrial 12S rRNA,16S rRNA and PreRNA 12S-16S in sucrose gradients using qRT-PCR.

To further investigate the assembly of ribosomal subunits, mitochondria were extracted from AWBSCR16 KO iBAT, dissolved, and separated with 10%–30% sucrose gradients. We observed a significant shift of the components in the *Wbscr16^-/-^* large ribosomal subunit to higher-density sucrose fractions, which differed from the Floxed ribosome (Fig. 2F). This indicates that the assembly of large ribosomal subunits in AWBSCR16 KO iBAT might be severely impaired. Furthermore, the distribution of mitochondrial transcripts within the ribosomal fractions was examined. In *Wbscr16* knockout mice, both 12S and 16S rRNA shifted to lower density fractions of the sucrose gradient, which co-migrated with the unprocessed polycistronic transcripts that contain both rRNAs (Fig. 2G-H). These findings confirmed that WBSCR16 knockout led to the disassembly of large ribosomal subunits in mitochondria and the inhibition of translational activities of mitochondria-encoded proteins.

### Requirement of WBSCR16 for 16S rRNA-tRNA^Leu^ Cleavage

To investigate the possible mechanism in the regulation of 16S rRNA, we performed RNA immunoprecipitation of FLAG-tagged WBSCR16 in MEFs followed by RIP-Seq and qRT-PCR. Referring to binding site sequences identified here, 16S rRNA is the only highly enrichment transcript associated with WBSCR16 across the entire mitochondrial transcriptome (Fig. 3A-B), consistent with the above data that only 16S rRNA dramatically reduced in AWBSCR16 KO iBAT (Fig. 2C). Consistently, it was further confirmed by Northern blot assay, which showed a marked reduction in mature 16S rRNA levels in AWBSCR16 KO tissues, while 12S rRNA levels remained unchanged (Fig. 3C). 16S and 12S rRNA are encoded by the initiation region of the mitochondrial heavy strand DNA ^31^. As shown in Fig. 3D, 12S rRNA is located at the 5’-end of 16S rRNA, while tRNA^Val^, tRNA^Leu^, and Nd1 are located at the 3’-end of 16S rRNA. We found significant enrichment of unprocessed transcripts containing tRNA within the 12S and 16S fragments in AWBSCR16 KO adipocytes, with a particularly notable increase in 16S rRNA-tRNA^Leu^-Nd1 transcripts (Fig. 3E). To further analyze the transcripts in BAT, we performed transcriptome-wide RNA sequencing (RNA-seq) of iBAT. The RNA-seq data confirmed the significant decrease in 16S rRNA levels and revealed an enrichment of reads spanning tRNA regions between 12S and 16S, as well as between 16S and Nd1, Co3 and Nd3, and Nd3 and Nd4L, indicating an accumulation of precursor RNAs (Fig. 3F). Further analysis revealed a pronounced decrease in cleavage at the 3’-end of 16S rRNA, which corresponds to the 5’-end of mt-tRNA^Leu^ (Fig. 3G). Thus, the mitochondrial changes observed in Fig. 1 were attributed to the loss of WBSCR16, which resulted in aberrant RNA splicing of 16S rRNA.

**Fig. 3.**
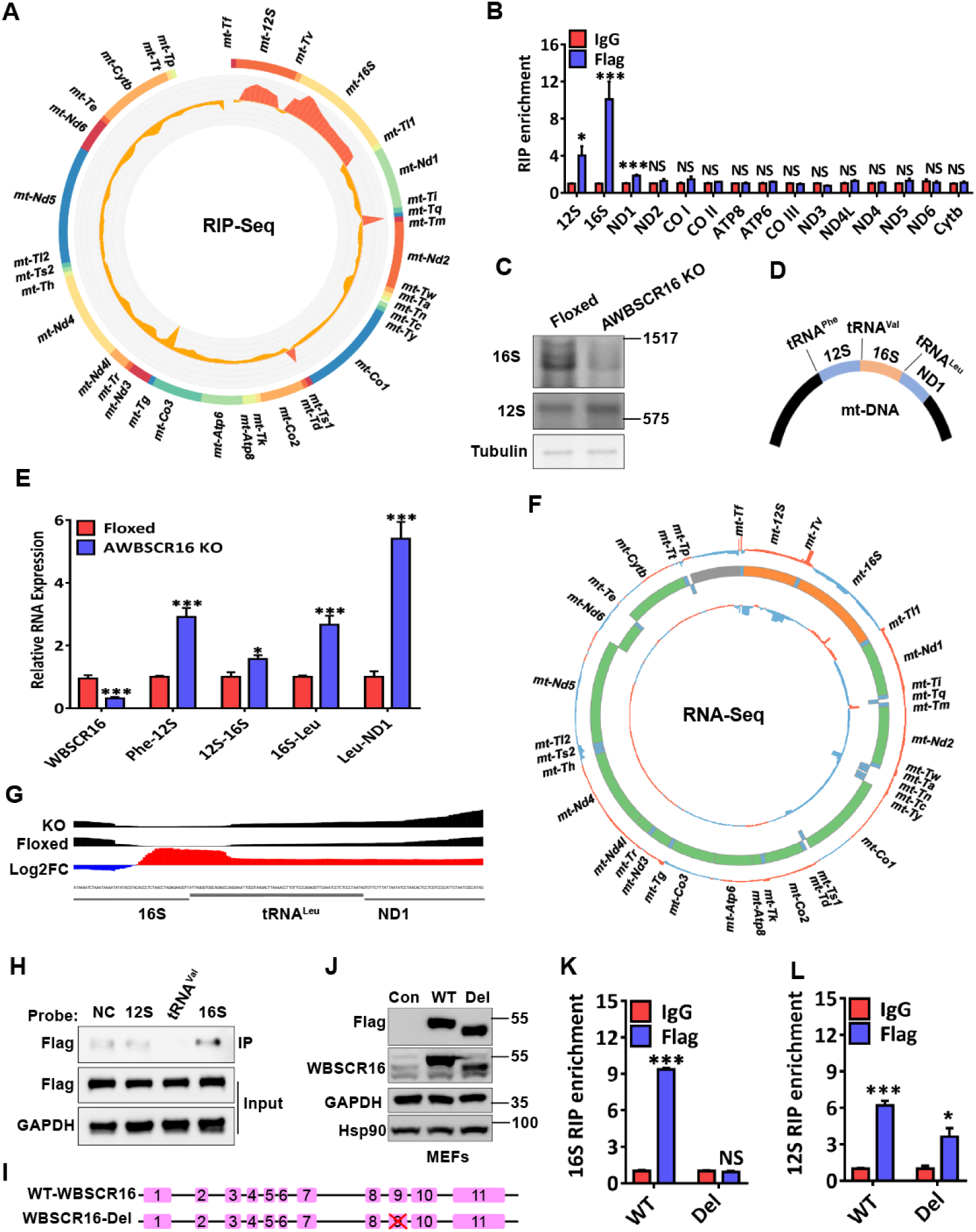
Mitochondrial 16S rRNA processing defects in AWBSCR16 KO cells and the requirement of the 9^th^ exon for 16S rRNA-WBSCR16 binding. **(A)** Analysis of the enrichment sequence of FLAG-tagged WBSCR16 binding using RIP-Seq. **(B)** qRT-PCR analysis of WBSCR16 and mtDNA-encoded RNAs in iBAT from Floxed and AWBSCR16 KO mice. **(C)** Northern blot analysis of 16S and 12S rRNA in Floxed and AWBSCR16 KO iBAT. Tubulin mRNA was detected as loading control. **(D)** Schematic illustration of the mitochondrial DNA fragment encoding rRNA (12S and 16S) and mRNA ORF punctuated by tRNAs. **(E)** Analysis of mitochondrial precursor RNA transcripts involved in 16S rRNA in Floxed and AWBSCR16 KO iBAT using qRT-PCR. **(F)** Complete map showing changes in RNA-seq coverage (log2 fold change [KO_mean_/Floxed_mean_]) from three control (Floxed) and three knockout (KO) mice, with increases shown in red and decreases in blue. The mitochondrial genome is displayed, with rRNAs in orange, mRNAs in green, tRNAs in blue, and the non-coding region (NCR) in gray. **(G)** Genome browser view showing the mean RNA-seq coverage (top: the raw obtained sequencing reads of three control (Floxed) and three knockout (KO) mice; bottom: log2 fold change [KO_mean_/Floxed_mean_]) displaying the 5’cleavage sites of mt-tRNA^Leu^ in the absence of WBSCR16. **(H)** Immunoblotting analysis of the pull-down products from FLAG-tagged WBSCR16 expressed in MEFs with biotin-labeled 12S rRNA, tRNA^Val^ and 16S rRNA specific DNA probes. **(I)** Schematic illustrated the exons in WT-WBSCR16 (WT) or Del-WBSCR16 (Del) protein. **(J)** Representative western blots of Flag and WBSCR16 in WT- or Del-WBSCR16 overexpression MEFs. **(K-L)** Immunoprecipitation of FLAG-tagged WT- and Del-WBSCR16 expressed in MEFs by FLAG beads, followed by qRT-PCR analysis of 16S rRNA(**K**) and 12S rRNA (**L**), normalized to GAPDH.

### WBSCR16 binds to 16S rRNA through a specific RNA-binding domain (RBD)

To further investigate the specific involvement of WBSCR16 in the maturation process of 16S rRNA, RNA pull-down experiments using DNA probes targeting 16S rRNA confirmed the specific binding of WBSCR16 to 16S rRNA transcripts (Fig. 3H). Our previous data showed that the mouse WBSCR16 gene consists of 11 exons, and the deletion of the 9th exon (Del-WBSCR16) in WBSCR16 leads to lethality in mice^24^. The deleted region localizes in the C-D loop between B6 and B7 folds, which is a potential RBD domain based on the crystallization structure ^32^. To determine which domains of WBSCR16 are involved in the interaction between 16S rRNA and WBSCR16, we generated stable overexpression of the 9th exon deletion mutant of WBSCR16 in MEFs (Fig. 3I-J). The RNA immunoprecipitation assay revealed that the mutant WBSCR16 failed to interact with 16S rRNA (Fig. 3K). This suggests that the deleted domain may contain the RBD within WBSCR16, which is crucial for its interaction with 16S rRNA, rather than 12S rRNA (Fig. 3L).

### WBSCR16 Interaction with MRPP3 *in vitro* and *in vivo*

RNase P and RNase Z are responsible for general RNA processing specifically in mitochondria ^17, 18^. We further investigated whether WBSCR16 regulates 16S rRNA through either of these enzymes. Interestingly, the expression of MRPP3, the catalytic subunit of RNase P, showed opposite fluctuation upon WBSCR16 knockout or overexpression. However, no changes were found for ELAC2 and other proteins involved in regulating mitochondrial gene expression, such like FASTKD2, MTERFD1, RPUSD4 and TRUB2 (Fig. S2A-H). The alterations of MRPP3 expression after WBSCR16 knockout or overexpression in BAT suggested a possible close relationship between these two proteins. To explore the possible interaction between MRPP3 and WBSCR16, immunoprecipitation experiments were conducted using WBSCR16-overexpression BAT or MEFs extracts, which revealed co-precipitation of MRPP3 with WBSCR16 and vice versa, but not ELAC2 (Fig. 4A-B). Similar results were observed in other tissues, such as the liver (Fig. S2I). Additionally, WBSCR16-Del, lacking the exon 9 domain, failed to interact with MRPP3 when precipitated from WT-WBSCR16 and WBSCR16-Del-overexpressing MEFs and BAT (Fig. 4C-D). This indicates that the exon 9 domain is critical for the interaction of WBSCR16 with both MRPP3 and 16S rRNA. To further investigate the *in vitro* interaction between WBSCR16 and MRPP3, pull-down assays were performed using recombinant WBSCR16-GST and MRPP3-His proteins, which confirmed a direct interaction between WBSCR16 and MRPP3 (Fig. 4E-F). These results are consistent with the possibility that these two proteins form complexes *in vivo*.

**Fig. 4.**
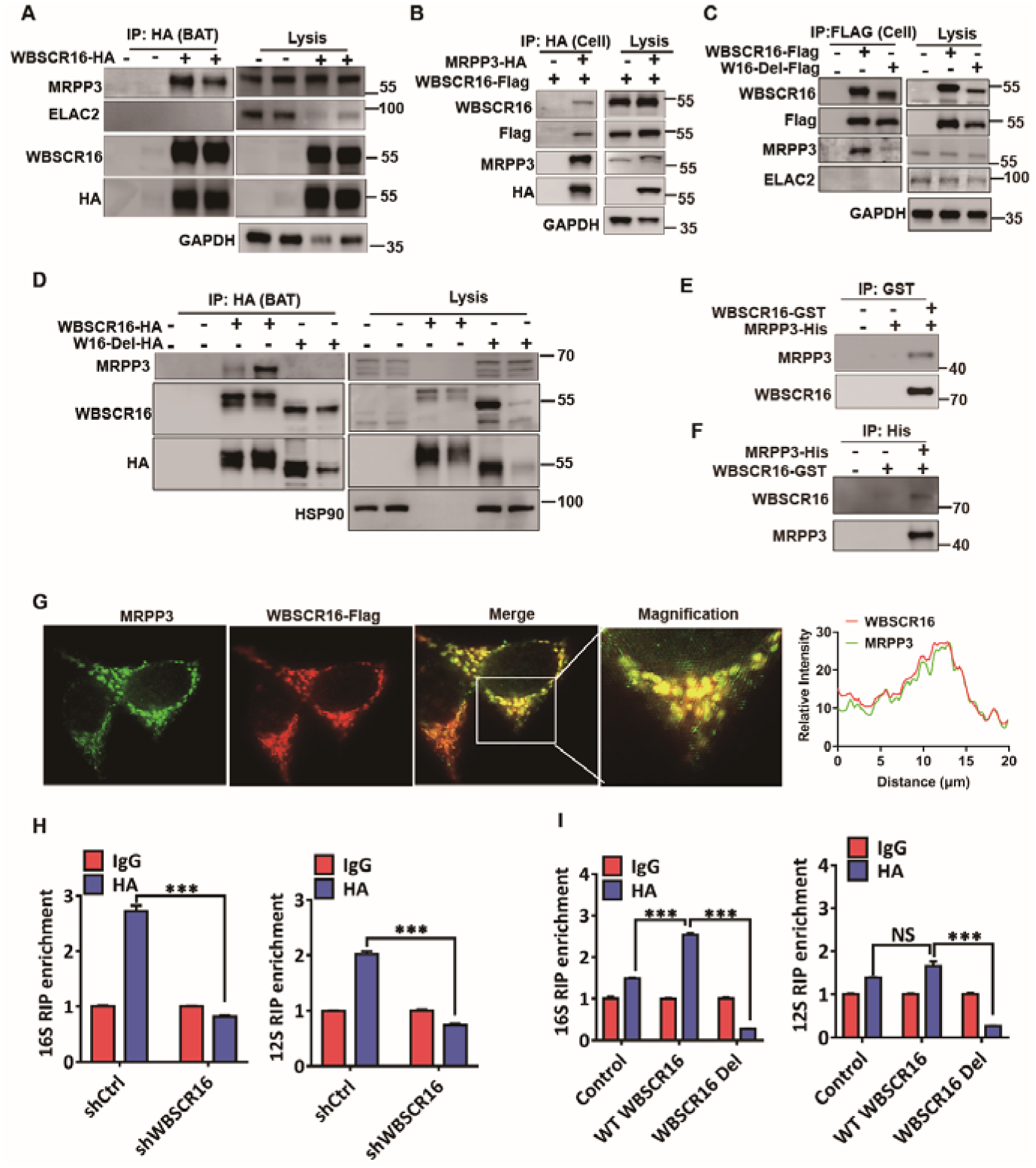
WBSCR16 interaction and colocalization with MRPP3, contributing to the binding ability between MRPP3 and 16S or 12S rRNA. **(A)** WBSCR16-HA overexpression BAT lysate immunoprecipitated with anti-HA beads, followed by immunoblotting with indicated antibodies. **(B)** MEF cells expressing with WBSCR16-Flag and MRPP3-HA, immunoprecipitated with anti-HA, followed by immunoblotting with indicated antibodies. **(C)** Immunoprecipitated of WBSCR16-Flag or WBSCR16-Del-Flag overexpressed cell lysate with anti-Flag with, followed by immunoblotted with indicated antibodies. **(D)** Co-immunoprecipitation of MRPP3 in lysed WBSCR16-Flag or WBSCR16-Del-Flag overexpressed BAT tissues. Immunoprecipitated with anti-HA bead, followed by immunoblotting with indicated antibodies. **(E-F)** *In vitro* interaction between WBSCR16 and MRPP3 analyzed with GST- (**E**) or His-(**F**) pull-down assays. **(G)** Immunofluorescent co-localization of WBSCR16 and MRPP3 in MEF cells. **(H-I)** RNA-immunoprecipitation of MRPP3 performed with an anti-HA antibody or anti-IgG antibodies in MRPP3-HA overexpression and shCtrl or shWBSCR16 (**H**), and in WT-WBSCR16-Flag or WBSCR16-Del-Flag overexpression in MEF cells (**I**), followed by qRT-PCR for 12S and 16S levels, which normalized to GAPDH.

To determine whether WBSCR16 and MRPP3 co-localize *in vivo*, immunofluorescent staining was performed on WBSCR16 overexpression cells, and we found the signal for WBSCR16 co-localized with MRPP3 in mitochondria (Fig. 4G). These results, along with the binding of WBSCR16 to 16S rRNA, support the potential formation of 16S rRNA:WBSCR16:MRPP3 tripartite complexes that cooperatively promote the processing of 16S rRNA. Based on these data, it is demonstrated that WBSCR16 can specifically bind to 16S rRNA and interact with MRPP3 to modify the 5’ site of mt-tRNA^Leu^ cleavage by RNase P, which in turn determines the 3’ processing of 16S rRNA.

To examine the involvement of WBSCR16 in MRPP3-mediated cleavage of 16S rRNA, we first established WBSCR16 knockdown in MRPP3 overexpressing cells. We then employed an RNA-immunoprecipitation assay to examine the effect of WBSCR16 knockdown on MRPP3 binding to mitochondrial-encoded RNA. The results showed a significant reduction in both 16S and 12S rRNA immunoprecipitated by MRPP3 when WBSCR16 was knocked down (Fig. 4H), while other mitochondrial-encoded genes were not affected (Fig. S3A-B). Furthermore, we also found that MRPP3 was able to bind more 16S rRNA and 12S rRNA when WT-WBSCR16 was overexpressed, but not for other mitochondrial RNAs (Fig. S3C-D). In contrast, the ability of MRPP3 to bind 16S rRNA and 12S rRNA was significantly reduced in WBSCR16-Del overexpressed cells, similar to WBSCR16 knockdown (Fig. 4I). These results suggest that WBSCR16 can directly bind to 16S rRNA, and also greatly enhance the ability of MRPP3 to bind to 16S rRNA and 12S rRNA. It is worth noting that while WBSCR16 assists MRPP3 in binding to both 16S rRNA and 12S rRNA, the main effect is observed in the processing of 16S rRNA due to its specific binding affinity for 16S rRNA, as demonstrated in Fig. 2. Additionally, the partial effect of WBSCR16 on the maturation of 12S rRNA can be attributed to its assistance in MRPP3 binding to 12S RNA (Fig. 3).

### Fat Utilization Preference in *Wbscr16^-/-^* BAT

To meet the high energy demands, metabolically active tissues are equipped with complex enzymatic machinery that efficiently orchestrates ATP production using multiple energy substrates, including FA, carbohydrates (glucose and lactate), ketones, and amino acids ^33–35^. Glucose oxidation and β-oxidation of FA occur within mitochondrial matrix, and the metabolic substrates in BAT fluctuate upon mitochondrial OXPHOS alteration ^36, 37^. Given the significant changes observed in Wbscr16-/- mitochondria, we assessed energy metabolism. Surprisingly, AWBSCR16 KO mice have higher plasma glucose but lower NEFA levels (Fig. 5A-B). To determine these differences, RNA-seq was then performed, which revealed upregulation of FA oxidation related genes, such as *Acot2, Scd3, Gpat3* and *Mogat*, among the most differentially expressed genes (DEGs) related to energy expenditure in WBSCR16 KO mice (Fig. 5C). Additionally, the KEGG pathway analysis of RNA-Seq data showed significant changes in lipid metabolism activity in AWBSCR16 KO mice (Fig. 5D). Furthermore, glucose metabolism shifting from glycolysis to the citrate cycle in matured adipocytes derived from iBAT-stromal vascular fraction (SVF) lacking WBSCR16 (Fig. S4A). Collectively, these data suggest that glucose metabolism activity decreases due to mitochondrial changes in AWBSCR16 KO iBAT, leading to a preference for lipid utilization as an energy source compared to Floxed mice.

**Fig. 5.**
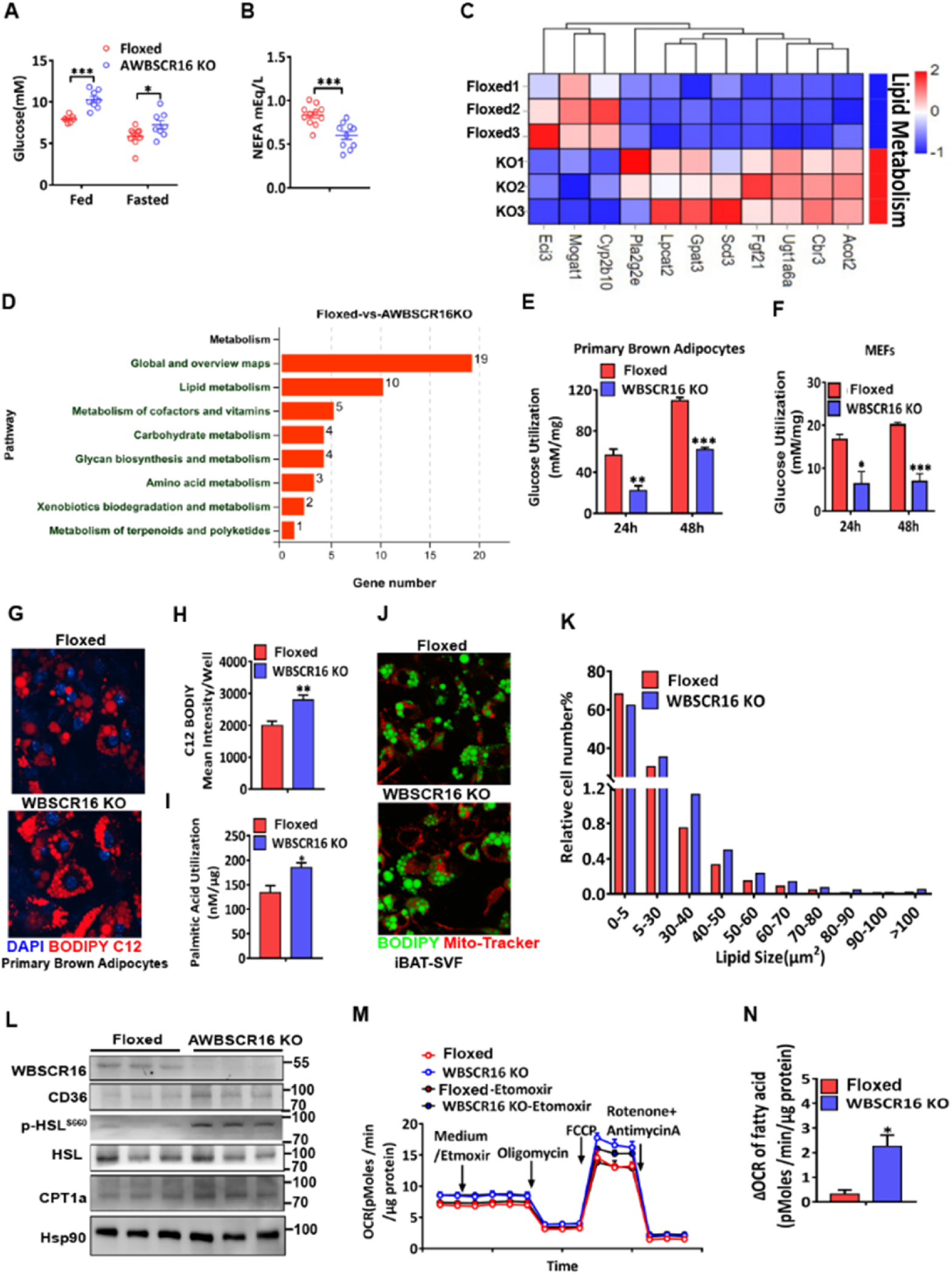
Metabolic flexibility by energy substrates switching in WBSCR16 deficient BAT. **(A-B)** Levels of fed and fasted glucose (**A**) and plasma NEFA (**B**) in Floxed and AWBSCR16 KO mice. **(C)** Heatmap displaying mRNA expression levels of upregulated or downregulated genes in iBAT from Floxed and AWBSCR16 KO mice. **(D)** KEGG pathway analysis of RNA-Seq from Floxed and AWBSCR16 KO iBAT. **(E-F)** Glucose utilization in matured adipocytes derived from Floxed iBAT-SVF infected with or without Cre lentivirus (**E**) or MEFs with WBSCR16 knockout (**F**) within 24 and 48 hours. Values were normalized by total protein. **(G-H)** Representative staining (**G**) and quantitative analysis (**H**) of the BODIPY C12 accumulated in lipid droplets of matured adipocytes derived from iBAT-SVF infected with or without Cre lentivirus. **(I)** Quantitative analysis of palmitic acid utilization in matured adipocytes derived from iBAT-SVF infected with or without Cre lentivirus within 24 hours. **(J-K)** Representative staining of lipid droplets and mitochondria with BODIPY and Mito-tracker, respectively (**J**), and quantitative analysis of lipid droplet size (**K**) in matured adipocytes derived from iBAT-SVF infected with or without Cre lentivirus. **(L)** Representative Western blots of CD36, p-HSL^S660^, total-HSL, CPT1α in Floxed and AWBSCR16 KO iBAT. **(M-N)** Substrate oxidation stress test of FA in matured adipocytes derived from iBAT-SVF infected with or without Cre lentivirus **(M)**. OCR came from FA after FCCP treatment was shown **(N)**.

To further confirm this assumption, glucose utilization in primary BAT and MEFs was detected. We found a significant decrease in glucose utilization in *Wbscr16^-/-^* cells after 24 or 48 hours (Fig. 5E-F). However, the ATP levels in these cells remained well-maintained, possibly due to compensatory FA oxidation (Fig. S4B-C). To further detect the rate of FA uptake, matured primary BAT cells were labeled with BODIPY C12 (Red C12), a FA-conjugated fluorescent probe. Quantitative analysis showed that *Wbscr16^-/-^* cells took up more FA than WT BAT cells (Fig. 5G-H). Moreover, when cultured with palmitate, *Wbscr16^-/-^* cells also used much more palmitate than Floxed BAT cells (Fig. 5I). Unimpaired adipogenesis and similar lipid size changes were observed in matured adipocytes derived from iBAT-SVF, further confirming these findings (Fig. 5J-K). However, no significant lipid changes were observed in primary WAT and iWAT-SVF, which have lower mitochondrial content (Fig. S4D-G). These results demonstrate that *Wbscr16^-/-^* cells switch their substrate preference from glucose to FA, contributing to the higher plasma glucose levels but lower NEFA levels in AWBSCR16 KO mice (Fig. 5A-B). This also explains the reduced triglyceride storage in *Wbscr16^-/-^* WAT.

Furthermore, proteins involved in regulating FA metabolism, such as CD36, HSL, and CPT1a, play a role in the transition from lipolysis to FA oxidation ^38–40^. Data showed a significant increase in CD36 expression, phosphorylation of HSL at Ser660, and CPT1a levels after WBSCR16 knockout tissue (Fig. 5L), indicating a shift towards prioritizing FA utilization in *Wbscr16^-/-^* adipocytes. Additionally, substrate-oxidation analysis was conducted to further confirm these findings, demonstrating that WBSCR16-deficient adipocytes preferentially utilize FA during oxygen consumption to produce ATP (Fig. 5M-N). As adipose tissue is an crucial regulator of various metabolic parameters ^41^, and thus, we assessed the effects on glucose tolerance and insulin resistance in AWBSCR16 knockout mice. Interestingly, AWBSCR16 knockout mice exhibited improved insulin resistance (Fig. S4H) despite have slightly higher plasma glucose levels and impaired glucose tolerance (Fig. 5A and Fig. S4I), compared to Floxed mice. These findings are consistent with previous studies indicating that decreased glucose uptake and increased reliance on FA oxidation are associated with impaired mitochondrial function ^42, 43^.

### Loss of WBSCR16 in Adipose Tissues Prevents Diet-Induced Obesity

To further determine the possible effects of adipose-specific WBSCR16 ablation on energy metabolism in mice, AWBSCR16 KO and Floxed mice were subjected to a high-fat diet (HFD) from 2 to 5 months of age. Interestingly, while there was no significant difference in total body weight between AWBSCR16 KO and Floxed mice on a regular chow diet (Fig. S1A), the control mice exhibited progressive weight gain on the HFD, whereas AWBSCR16 KO mice gained significantly less weight and maintained a relatively stable weight (Fig. 6A and Fig. S5A). By 5 months of age, AWBSCR16 KO mice under HFD were approximately 40% lighter than controls and had substantially reduced white adipose tissue (WAT) weight compared to control mice (Fig. 6B). The body weight of AWBSCR16 KO mice fed with high-fat diet remained similar to that of chow-diet fed mice (Fig. S5B), while the weight of high-fat diet Floxed mice increased markedly (Supplemental Fig. 5A). Furthermore, although there was a slight increase in the weight and lipid droplet size of iBAT in AWBSCR16 KO mice after HFD feeding, it was comparable to that of the chow diet (Fig. 6C). Importantly, AWBSCR16 KO mice had dramatically smaller iWAT and eWAT, which were only 20-25% of those in Floxed mice (Fig. 6D-E, 6G-H). Furthermore, there was no evidence of adipocyte atrophy in AWBSCR16 KO WAT, as the size of lipid droplets in WAT remained normal (Fig. 6D-E). The reduced size of AWBSCR16 KO WAT suggested lower triglyceride storage and increased lipolysis, with a higher utilization of fatty acids by iBAT. Although there were no differences in liver weight, plasma insulin levels, and glucose intolerance, there was a notable increase in lipid droplets in the liver of Floxed mice (Fig. 6F, Supplemental Fig. 5C-D). In contrast, AWBSCR16 KO mice maintained healthy WAT and liver, exhibited slightly higher plasma glucose levels, but lower plasma NEFA levels after 3 months HFD feeding (Fig. 6F-J), which was similar to chow-diet fed mice. Interestingly, despite the presence of mitochondrial dysfunction in BAT due to abnormal mitochondrial genomic expression, lipid metabolism was enhanced in AWBSCR16 KO mice (Fig. 6K), indicating the importance of WBSCR16-mediated mitochondrial changes in substrate preference and metabolic flexibility in mice. To investigate the potential correlation between WBSCR16 expression levels and human obesity, we examined microarray data from GSE27951, which includes expression profiles of subcutaneous adipose tissue obtained from 20 individuals with a range of body mass indices ^44^. Interestingly, the expression of WBSCR16 was found to be higher in obesity compared to non-obese individuals (Fig. 6L), suggesting a potential role for WBSCR16 in energy metabolism in humans.

**Fig. 6.**
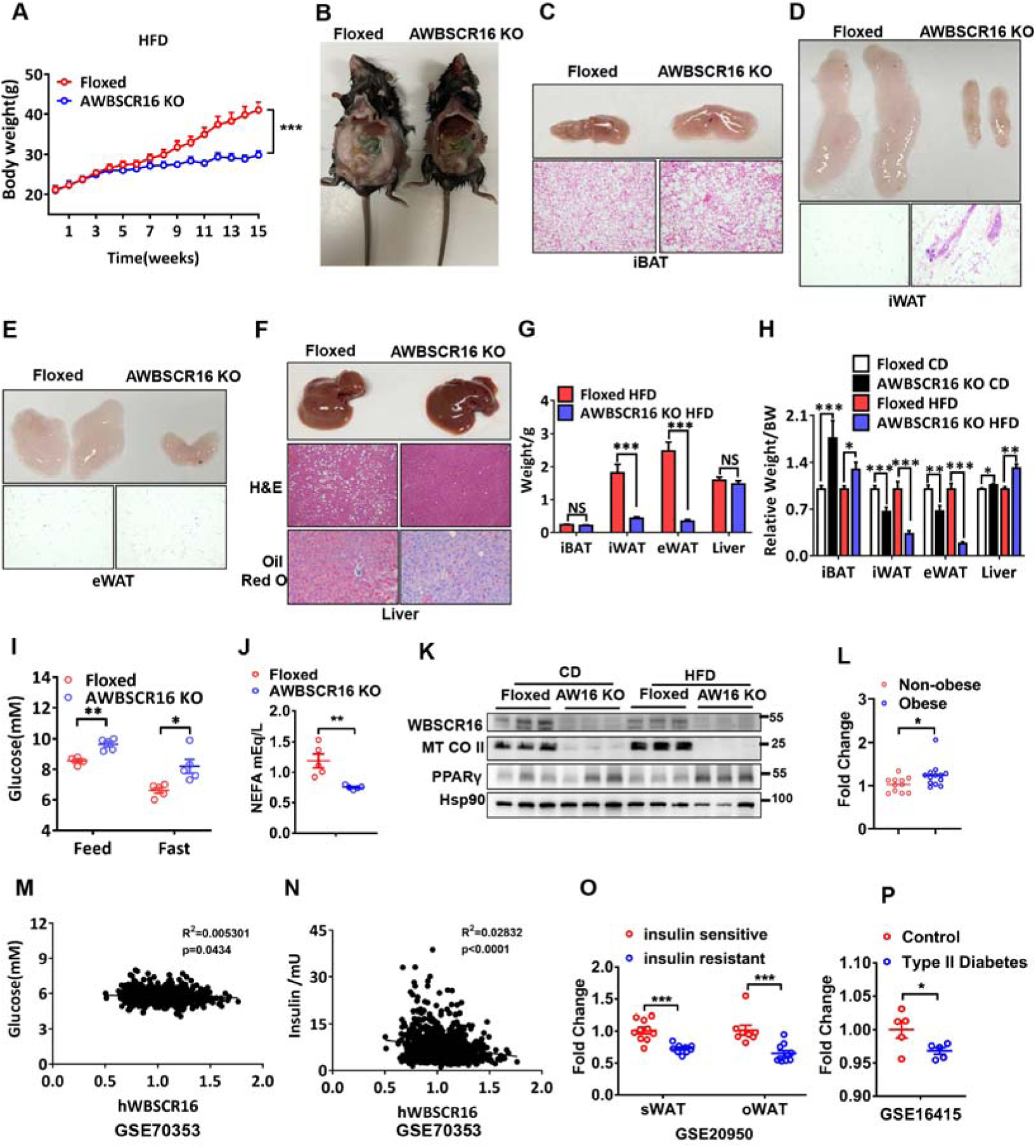
Deletion of WBSCR16 in Adipose Tissues Prevents Diet-induced Obesity. **(A)** Growth curve and representative picture **(B)** of HFD-fed Floxed and AWBSCR16 KO mice. **(C-E)** Representative pictures and HE staining of iBAT (**C**), iWAT (**D**) and eWAT (**E**) from mice after being HFD-fed. **(F)** Representative pictures, HE staining and Oil Red O staining of liver from HFD-fed mice. **(G)** Weights of iBAT, iWAT, eWAT, liver and their relative weights normalized to body weight (**H**) from HFD-fed mice. (**I**) Levels of fed and fasted glucose, fasted plasma NEFA **(J)** in HFD-fed Floxed and AWBSCR16 KO mice **(K)** Representative Western blots of WBSCR16, MT-CO II, and PPARγ in CD-fed or HFD-fed iBAT. **(L)** Analysis of WBSCR16 mRNA expression in adipose tissue from obese and non-obese subjects (BMI 16.7-50.2) with normal or impaired glucose tolerance (GSE27951).

### Overexpression of WBSCR16 in BAT

To further investigate the possible effects of WBSCR16 in mitochondria in BAT, transgenic mice with inducible WBSCR16 expression in BAT (WBSCR16^UCP1tg^) were obtained using the Cre-LoxP system (WBSCR16^tg/+^; UCP1 ERT-Cre^+/–^). After being induced, WBSCR16 expression was greatly increased in BAT (Fig. 7A). In contrast to AWBSCR16 KO mice, WBSCR16^UCP1tg^ mice exhibited lower plasma glucose levels (Fig. 7B), improved glucose tolerance (Fig. 7C-D), but impaired insulin sensitivity (Fig. S6A-B), attributed to increased glucose utilization in WBSCR16^UCP1tg^ mice. Additionally, WBSCR16 overexpression in MEFs also resulted in a preference for glucose usage (Fig. 7E). Interestingly, both the weight and lipid droplet size of transgenic iBAT were decreased (Fig. 7F-H), while the overall body weight, WAT weight, and fat content remained normal (Fig. S6C-G). These findings suggest that substrate preference and mitochondrial function in BAT play crucial roles in glucose tolerance and insulin resistance. Moreover, increased glucose utilization through WBSCR16 overexpression may help improve glucose intolerance in diabetes. More importantly, WBSCR16 transgenic iBAT displayed noticeable changes in mitochondrial morphology, including a significant increase in tubular cristae, slightly higher mitochondrial number, smaller mitochondrial size, and increased expression of mtDNA-encoded proteins (Fig. 7I-L). As anticipated, real-time PCR and Northern blot assays revealed increased expression of two mitochondrial ribosomal RNAs (12S and 16S), while unprocessed transcripts in WBSCR16 transgenic iBAT markedly decreased (Fig. 7M-O). These alterations in mitochondrial morphology and biogenesis were opposite to those observed in WBSCR16 knockout iBAT, indicating that WBSCR16 plays a significant role in mitochondrial OXPHOS functions. However, when subjected to a HFD for 3 months, WBSCR16 transgenic mice gained more body weight and fat mass compared to control mice (Fig. S6I-J), without affecting lipid deposition in the liver (Fig. S6K-L). These findings contrast with the phenotypes observed in AWBSCR16 KO mice fed a HFD (Fig. 6).

**Fig. 7.**
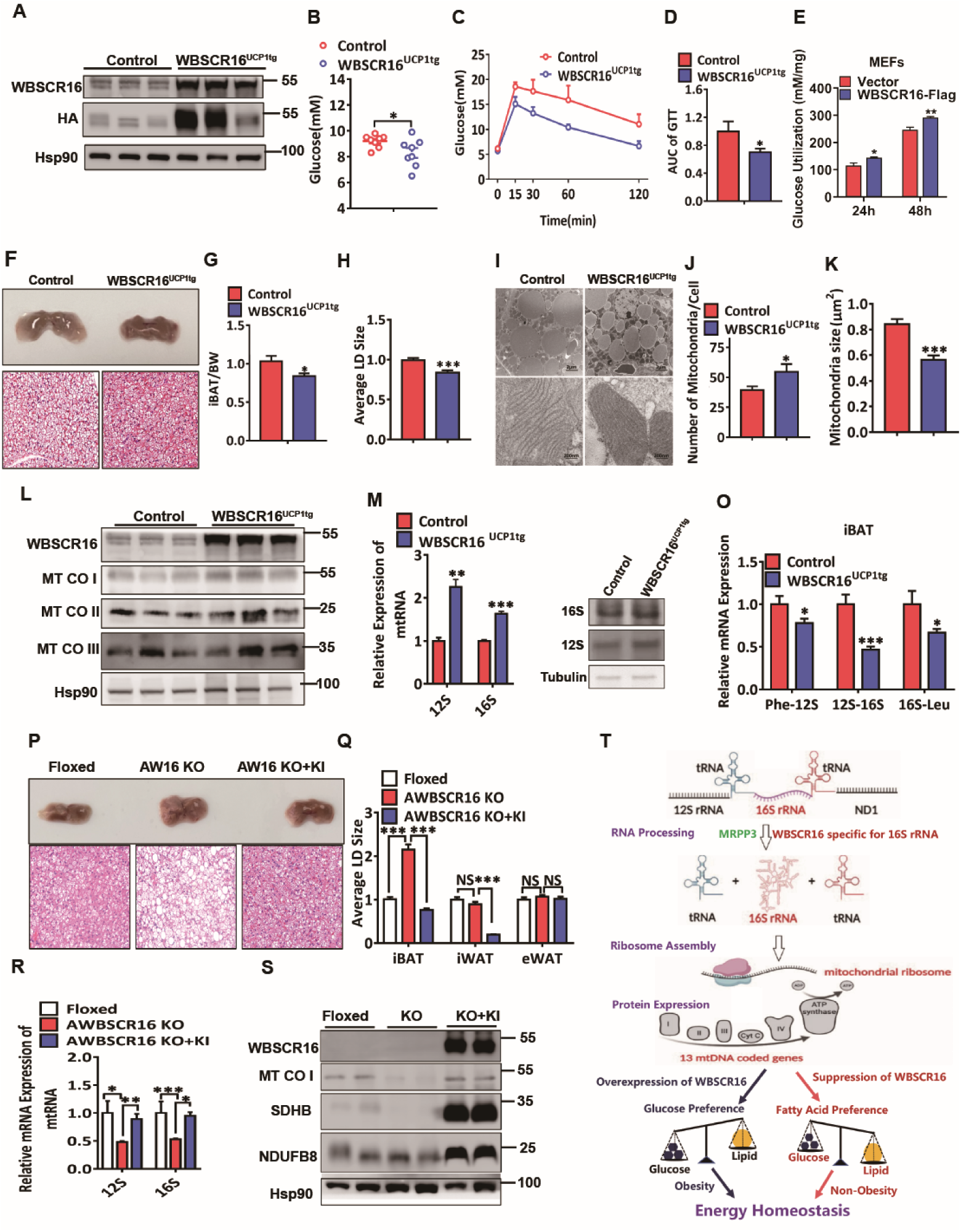
Overexpression of WBSCR16 improved mitochondria and exogenous WBSCR16 rescued AWBSCR16 KO in BAT. **(A)** Representative Western blots of WBSCR16 in iBAT from Floxed and WBSCR16^UCP1tg^ mice as indicated. **(B-D)** Resting glucose levels **(B)**, glucose tolerance test **(C),** and AUC analysis **(D)** of Floxed and WBSCR16^UCP1tg^ mice. **(E)** Glucose utilization in WBSCR16 overexpression MEFs. **(F-H)** Representative pictures and HE staining of iBAT from Control and WBSCR16^UCP1tg^ mice **(F)**. Relative weight normalized to body weight **(G)** and cross-sectional area of lipid droplets of iBAT (**H**) were shown. **(I-K)** Electron microscopy images of iBAT (**I**), number (**J**), size (**K**) of mitochondria from Control and WBSCR16^UCP1tg^ mice. **(L)** Representative Western blots of WBSCR16, MT-CO I, MT-CO II and MT-CO III in iBAT from Control and WBSCR16^UCP1tg^ mice as indicated. **(M)** qRT-PCR analysis of 12S and 16S rRNA in iBAT from Control and WBSCR16^UCP1tg^ mice. **(N)** Northern blot analysis of 16S and 12S rRNA in iBAT. mRNA of Tubulin was detected as a loading control. **(O)** Mitochondrial precursor RNA transcripts involved 16S rRNA were analyzed by qRT-PCR. **(P-Q)** Representative pictures and HE staining **(P)** and the size of lipid droplets **(Q)** in Floxed, AWBSCR16 KO and AWBSCR16 KO+KI iBAT. **(R)** qRT-PCR analysis of 12S and 16S in Floxed, AWBSCR16 KO, and AWBSCR16 KO+KI iBAT. **(S)** Representative Western blots of WBSCR16, MT-CO I, SDHB, NDUFB8 and HSP90 in iBAT as indicated. **(T)** Schematic diagram of the working model for the roles of WBSCR16 in mitochondria metabolic substrate preference *in vivo*.

### Rescued by Adipocyte-specific Exogenous WBSCR16 Completely

To validate whether the observed phenotype in AWBSCR16 KO mice is specifically attributed to WBSCR16 deletion, we generated WBSCR16^f/f^:WBSCR16^Tg/+^:Adipo Cre^+/-^ mice by crossing AWBSCR16 KO with AWBSCR16 KI mice. Remarkably, the overexpression of exogenous WBSCR16 completely restored the decreased levels of mature mitochondrial 16S rRNA and 12S rRNA, as well as the increase in immature ones caused by WBSCR16 deletion specifically, without altering other mtDNA-coded genes (Fig. 7R, Fig. S7A-B). Additionally, all the phenotypes observed in AWBSCR16 KO mice, such as larger lipid droplets in BAT and smaller ones in subcutaneous WAT, were fully rescued by exogenous WBSCR16 (Fig. 7Q-R and Fig. S7C-D). These findings indicate that the overexpression of exogenous WBSCR16 in specific tissues can enhance mitochondrial structural stability and alter substrate preference, thereby improving certain functions associated with metabolic diseases caused by defects in mitochondrial OXPHOS.

In summary, as depicted in the model presented in Fig. 7T, we identified mitochondrial 16S rRNA processing defects that contribute to mitochondrial dysfunction in *Wbscr16^-/-^* cells. Further investigations revealed that WBSCR16 functions as a 16S rRNA-binding protein and collaborates with MRPP3 to play essential roles in 16S rRNA processing. This disruption affects the assembly of the large mitochondrial ribosome, leading to reduced expression of mitochondrial proteins and impaired normal OXPHOS activities through the electron transport chain (ETCs) in mitochondria. Consequently, the fluctuating fuel utilization in *Wbscr16^-/-^* BAT results in abnormal whole-body energy expenditure and prevents weight gain following high-fat diet treatment. However, due to mitochondrial changes associated with WBSCR16 overexpression, cells exhibit a preference for glucose as an energy source, increasing susceptibility to obesity *in vivo* (Fig. 7T).

## Discussion

In this study, we investigated the role of WBSCR16 in adipose tissues and its impact on energy metabolism. Our findings highlight the importance of WBSCR16 in mitochondrial function and substrate preference within BAT. We demonstrated that WBSCR16 is involved in the processing of mitochondrial 16S rRNA and its deletion leads to mitochondrial dysfunction, disrupted OXPHOS, and impaired glucose and lipid metabolism in BAT.

Fat in the body exhibits different colors that correspond to its function, BAT is a key regulator of energy expenditure and whole-body homeostasis, primarily through FA oxidation or thermogenesis ^1, 2^. The metabolic substrates in BAT, including glucose and fatty acids, fluctuate depending on mitochondrial OXPHOS activity ^36, 37^. Although blocking the utilization of these substrates suppress BAT activation significantly ^45^, specific functions of them in BAT is unclear and requires further study. In our study, the loss of WBSCR16 resulted in decreased OXPHOS in BAT, leading to a dramatic impairment in glucose metabolism and tricarboxylic acid cycle. The disassembly of the large ribosomal subunit and inhibition of protein synthesis within mitochondria further contributed to the metabolic alterations. These findings emphasize the crucial role of mitochondrial glucose/lipid metabolism in BAT for maintaining whole-body energy homeostasis.

Interestingly, some phenotypes observed in the AWBSCR16 KO mice are similar to the characteristics observed in mice fed a ketogenic diet, such as reduced weight gain and increased FA utilization ^46, 47^. However, there are notable differences between AWBSCR16 KO mice and mice on a ketogenic diet. AWBSCR16 KO mice exhibit mildly elevated fasting glucose levels, decreased free fatty acids (FFA), and unchanged insulin levels in their plasma, whereas mice on a ketogenic diet primarily rely on fatty acids as their main energy substrate within the mitochondria, resulting in lower plasma glucose and insulin levels similar to those observed during starvation ^48^. Despite these distinctions, the AWBSCR16 KO mice consistently demonstrate decreased FFA levels, whether they are on a chow diet or a high-fat diet, leading to smaller white adipose tissue (WAT) depots and reduced fat accumulation in the liver. The compensatory increase in fatty acid utilization in brown adipose tissue (BAT) is particularly noteworthy, as it promotes whole-body energy expenditure and even prevents high-fat diet-induced obesity. In summary, while a ketogenic diet involves passive fat absorption and utilization, providing minimal carbohydrates but predominantly fat (over 80%) as energy substrates, AWBSCR16 KO cells and mice exhibit an active preference for fatty acid utilization regardless of the diet provided. Therefore, we propose a novel mechanism and potential target for reducing lipid accumulation and combating obesity resistance.

AWBSCR16 KO not only leads to decreased lipid levels in mice but also enhances insulin sensitivity, thereby potentially ameliorating diabetes resulting from insulin resistance (Fig. 5), even under high-fat fed (Fig. 6). Conversely, overexpression of WBSCR16 in mice promotes a preference for glucose utilization, significantly improving glucose tolerance and reducing blood sugar levels compared to the control group (Fig. 7). These findings suggest that different forms of diabetes may be improved by either overexpressing WBSCR16 (in cases of glucose intolerance) or inhibiting WBSCR16 (in cases of insulin resistance) in BAT via differential metabolic substrate preferences. Thus, the WBSCR16-mediated mitochondrial substrate preference plays a crucial role not only in the development and treatment of obesity but also in different forms of diabetes, offering valuable insights into the understanding of obesity and diabetes.

The absence of WBSCR16 results in a significant reduction in 16S rRNA levels, consequently inhibiting the assembly of the large mitochondrial ribosomal subunit. Similar changes have been observed in cells deficient in RNA-binding proteins (RBPs) such as PTCD1 or FASTKD2, which are known to regulate the stability of mt-Nd6 mRNA and 16S rRNA ^49, 50^. However, the specific roles of these RBPs in the processing of 16S rRNA remain unknown. In our study, we found that WBSCR16 directly binds to 16S rRNA and interacts with MRPP3, which is the first identified RBP to specifically promote the maturation of 16S rRNA. On the other hand, WBSCR16 deficiency in BAT leads to increased triglyceride accumulation, as evidenced by elevated levels of CD36, p-HSL(S660), and CPT1a, which aligns with previous findings in PTCD1 KO cells ^51^. Other studies suggest that decreased glucose uptake and increased reliance on fatty acid oxidation are early events associated with impaired mitochondrial function in conditions such as Type I diabetes and obesity ^42, 43^. The altered balance between glucose and fatty acid utilization in BAT observed in AWBSCR16 KO mice leads to abnormal whole-body energy expenditure and contributes to energy wastage. These findings provide insights into a novel mechanism and potential treatment approaches for insulin resistance, glucose intolerance, and metabolic flexibility.

## Methods

### Animals

All animal experiments were performed according to procedures approved by the Institutional Animal Care and Use Committee for animal care and handling at Shanghai Jiaotong University School of Medicine. WBSCR16 (RCC1L)^flox/flox^ mice, WBSCR16 (RCC1L)^tg/tg^ mice and Adiponectin-cre mice were obtained from the Cyagen Biosciences. mUcp1-CreERT2 mice and obesity human samples were gifts from Professor Jiqiu Wang in Shanghai Institute for Endocrine and Metabolic Diseases. Mice were maintained on a standard rodent chow at ambient temperature (22℃) under 12 hours light-12 hours dark cycles. Littermates expressing no Cre (Floxed mice) were used as a control group within above animal experiments. Samples were obtained from male mice at 8-10 weeks.

### RNA-Seq and Alignments

RNA sequencing was performed on total RNA extrated from three Floxed and three AWBSCR16 KO mice after removing cytoplasmic rRNA().The RNA sequencing was carried out by using the Illumina HiSeq platform, according to the Illumina Tru-Seq protocol Library preparation, sequencing, and bioinformatics analyses.

### Statistical Analysis

All of the experiments were done at least in triplicate, or otherwise indicated. Data of Western Blot was digitalized and analyzed using either the Image J software or the histogram panel of Adobe Photoshop. Statistical analyses were performed using the Prism-7 software and Microsoft Excel. The data are presented as the mean + S.E.M. of absolute values or percentages of control. The values obtained *in vivo* or *in vitro* for the different parameters studied were compared using a Student’s two-tailed unpaired t-test for comparison of two groups. For comparison of multiple groups, we performed one-way analysis of variance (ANOVA) for all the groups of the experiment and showed as below: *p<0.05; **p<0.01; ***p<0.001; NS, not significant.

## Acknowledgments

This work was supported by the National Natural Science Foundation of China (81770845 and 91854124) (To Guorui Huang), (82088102) (To Guang Ning) and (32100923) (To Shengjie Zhang), National Natural Science Foundation of China (82070880) (To Zhiyun Zhao), Shanghai Science & Technology Foundation (18140902800) (To Guorui Huang).

## Author contributions

S.J.Z., Z.D., G.N. and G.R.H. designed the experiments, analyzed data, and wrote the manuscript; S.J.Z., Z.D., Y.F., W.G., S.Y.F., Z.Y.L. Z.Y.Z. N.C. and G.R.H carried out the experiments; S.J.Z., Z.D., G.N. and G.R.H. analyzed data and wrote the manuscript.

## References

1 Cypess, A. M. Reassessing Human Adipose Tissue. N Engl J Med 386, 768–779, doi:10.1056/NEJMra2032804 (2022).

2 Cohen, P. & Kajimura, S. The cellular and functional complexity of thermogenic fat. Nat Rev Mol Cell Biol 22, 393–409, doi:10.1038/s41580-021-00350-0 (2021).

3 Labbe, S. M. et al. In vivo measurement of energy substrate contribution to cold-induced brown adipose tissue thermogenesis. FASEB J 29, 2046–2058, doi:10.1096/fj.14-266247 (2015).

4 Iwen, K. A. et al. Cold-Induced Brown Adipose Tissue Activity Alters Plasma Fatty Acids and Improves Glucose Metabolism in Men. J Clin Endocrinol Metab 102, 4226–4234, doi:10.1210/jc.2017-01250 (2017).

5 Samuelson, I. & Vidal-Puig, A. Studying Brown Adipose Tissue in a Human in vitro Context. Front Endocrinol (Lausanne) 11, 629, doi:10.3389/fendo.2020.00629 (2020).

6 Osuna-Prieto, F. J. et al. Activation of Human Brown Adipose Tissue by Capsinoids, Catechins, Ephedrine, and Other Dietary Components: A Systematic Review. Adv Nutr 10, 291–302, doi:10.1093/advances/nmy067 (2019).

7 Storlien, L., Oakes, N. D. & Kelley, D. E. Metabolic flexibility. Proc Nutr Soc 63, 363–368, doi:10.1079/PNS2004349 (2004).

8 Galgani, J. E., Moro, C. & Ravussin, E. Metabolic flexibility and insulin resistance. Am J Physiol Endocrinol Metab 295, E1009–1017, doi:10.1152/ajpendo.90558.2008 (2008).

9 Fernandez-Verdejo, R., Marlatt, K. L., Ravussin, E. & Galgani, J. E. Contribution of brown adipose tissue to human energy metabolism. Mol Aspects Med 68, 82–89, doi:10.1016/j.mam.2019.07.003 (2019).

10 Yamaguchi, S. et al. Adipose tissue NAD(+) biosynthesis is required for regulating adaptive thermogenesis and whole-body energy homeostasis in mice. Proc Natl Acad Sci U S A 116, 23822–23828, doi:10.1073/pnas.1909917116 (2019).

11 Bodis, K. & Roden, M. Energy metabolism of white adipose tissue and insulin resistance in humans. Eur J Clin Invest 48, e13017, doi:10.1111/eci.13017 (2018).

12 Seyfried, T. N., Arismendi-Morillo, G., Mukherjee, P. & Chinopoulos, C. On the Origin of ATP Synthesis in Cancer. iScience 23, 101761, doi:10.1016/j.isci.2020.101761 (2020).

13 Rolfe, D. F. & Brown, G. C. Cellular energy utilization and molecular origin of standard metabolic rate in mammals. Physiol Rev 77, 731–758, doi:10.1152/physrev.1997.77.3.731 (1997).

14 Scarpulla, R. C. Nuclear control of respiratory chain expression by nuclear respiratory factors and PGC-1-related coactivator. Ann N Y Acad Sci 1147, 321–334, doi:10.1196/annals.1427.006 (2008).

15 Kobayashi, A., Azuma, K., Ikeda, K. & Inoue, S. Mechanisms Underlying the Regulation of Mitochondrial Respiratory Chain Complexes by Nuclear Steroid Receptors. Int J Mol Sci 21, doi:10.3390/ijms21186683 (2020).

16 Gustafsson, C. M., Falkenberg, M. & Larsson, N. G. Maintenance and Expression of Mammalian Mitochondrial DNA. Annu Rev Biochem 85, 133–160, doi:10.1146/annurev-biochem-060815-014402 (2016).

17 Holzmann, J. et al. RNase P without RNA: identification and functional reconstitution of the human mitochondrial tRNA processing enzyme. Cell 135, 462–474, doi:10.1016/j.cell.2008.09.013 (2008).

18 Siira, S. J. et al. Concerted regulation of mitochondrial and nuclear non-coding RNAs by a dual-targeted RNase Z. EMBO Rep 19, doi:10.15252/embr.201846198 (2018).

19 Takaku, H., Minagawa, A., Takagi, M. & Nashimoto, M. A candidate prostate cancer susceptibility gene encodes tRNA 3’ processing endoribonuclease. Nucleic Acids Res 31, 2272–2278, doi:10.1093/nar/gkg337 (2003).

20 Sanchez, M. I. et al. RNA processing in human mitochondria. Cell Cycle 10, 2904–2916, doi:10.4161/cc.10.17.17060 (2011).

21 Rackham, O. et al. Hierarchical RNA Processing Is Required for Mitochondrial Ribosome Assembly. Cell Rep 16, 1874–1890, doi:10.1016/j.celrep.2016.07.031 (2016).

22 Metodiev, M. D. et al. Methylation of 12S rRNA is necessary for in vivo stability of the small subunit of the mammalian mitochondrial ribosome. Cell Metab 9, 386–397, doi:10.1016/j.cmet.2009.03.001 (2009).

23 Metodiev, M. D. et al. NSUN4 is a dual function mitochondrial protein required for both methylation of 12S rRNA and coordination of mitoribosomal assembly. PLoS Genet 10, e1004110, doi:10.1371/journal.pgen.1004110 (2014).

24 Huang, G. et al. WBSCR16 Is a Guanine Nucleotide Exchange Factor Important for Mitochondrial Fusion. Cell Rep 20, 923–934, doi:10.1016/j.celrep.2017.06.090 (2017).

25 Reyes, A., Favia, P., Vidoni, S., Petruzzella, V. & Zeviani, M. RCC1L (WBSCR16) isoforms coordinate mitochondrial ribosome assembly through their interaction with GTPases. PLoS Genet 16, e1008923, doi:10.1371/journal.pgen.1008923 (2020).

26 Arroyo, J. D. et al. A Genome-wide CRISPR Death Screen Identifies Genes Essential for Oxidative Phosphorylation. Cell Metab 24, 875–885, doi:10.1016/j.cmet.2016.08.017 (2016).

27 Antonicka, H. et al. A pseudouridine synthase module is essential for mitochondrial protein synthesis and cell viability. EMBO Rep 18, 28–38, doi:10.15252/embr.201643391 (2017).

28 Jiao, X. et al. PHIP - a novel candidate breast cancer susceptibility locus on 6q14.1. Oncotarget 8, 102769–102782, doi:10.18632/oncotarget.21800 (2017).

29 Enerback, S. The origins of brown adipose tissue. N Engl J Med 360, 2021–2023, doi:10.1056/NEJMcibr0809610 (2009).

30 Cypess, A. M. & Kahn, C. R. Brown fat as a therapy for obesity and diabetes. Curr Opin Endocrinol Diabetes Obes 17, 143–149, doi:10.1097/MED.0b013e328337a81f (2010).

31 Montoya, J., Christianson, T., Levens, D., Rabinowitz, M. & Attardi, G. Identification of initiation sites for heavy-strand and light-strand transcription in human mitochondrial DNA. Proc Natl Acad Sci U S A 79, 7195–7199, doi:10.1073/pnas.79.23.7195 (1982).

32 Koyama, M., Sasaki, T., Sasaki, N. & Matsuura, Y. Crystal structure of human WBSCR16, an RCC1-like protein in mitochondria. Protein Sci 26, 1870–1877, doi:10.1002/pro.3210 (2017).

33 Alghannam, A. F., Ghaith, M. M. & Alhussain, M. H. Regulation of Energy Substrate Metabolism in Endurance Exercise. Int J Environ Res Public Health 18, doi:10.3390/ijerph18094963 (2021).

34 McNeill, B. T., Morton, N. M. & Stimson, R. H. Substrate Utilization by Brown Adipose Tissue: What’s Hot and What’s Not? Front Endocrinol (Lausanne) 11, 571659, doi:10.3389/fendo.2020.571659 (2020).

35 Jones, J. G. Hepatic glucose and lipid metabolism. Diabetologia 59, 1098–1103, doi:10.1007/s00125-016-3940-5 (2016).

36 De Meis, L., Ketzer, L. A., Camacho-Pereira, J. & Galina, A. Brown adipose tissue mitochondria: modulation by GDP and fatty acids depends on the respiratory substrates. Biosci Rep 32, 53–59, doi:10.1042/BSR20100144 (2012).

37 Vacanti, N. M. et al. Regulation of substrate utilization by the mitochondrial pyruvate carrier. Mol Cell 56, 425–435, doi:10.1016/j.molcel.2014.09.024 (2014).

38 Zhou, D. et al. CD36 level and trafficking are determinants of lipolysis in adipocytes. FASEB J 26, 4733–4742, doi:10.1096/fj.12-206862 (2012).

39 Anthonsen, M. W., Ronnstrand, L., Wernstedt, C., Degerman, E. & Holm, C. Identification of novel phosphorylation sites in hormone-sensitive lipase that are phosphorylated in response to isoproterenol and govern activation properties in vitro. J Biol Chem 273, 215–221, doi:10.1074/jbc.273.1.215 (1998).

40 Calderon-Dominguez, M. et al. Carnitine Palmitoyltransferase 1 Increases Lipolysis, UCP1 Protein Expression and Mitochondrial Activity in Brown Adipocytes. PLoS One 11, e0159399, doi:10.1371/journal.pone.0159399 (2016).

41 Rondinone, C. M. Adipocyte-derived hormones, cytokines, and mediators. Endocrine 29, 81–90, doi:10.1385/endo:29:1:81 (2006).

42 Buchanan, J. et al. Reduced cardiac efficiency and altered substrate metabolism precedes the onset of hyperglycemia and contractile dysfunction in two mouse models of insulin resistance and obesity. Endocrinology 146, 5341–5349, doi:10.1210/en.2005-0938 (2005).

43 Armoni, M., Harel, C., Bar-Yoseph, F., Milo, S. & Karnieli, E. Free fatty acids repress the GLUT4 gene expression in cardiac muscle via novel response elements. J Biol Chem 280, 34786–34795, doi:10.1074/jbc.M502740200 (2005).

44 Keller, P. et al. Gene-chip studies of adipogenesis-regulated microRNAs in mouse primary adipocytes and human obesity. BMC Endocr Disord 11, 7, doi:10.1186/1472-6823-11-7 (2011).

45 Veliova, M. et al. Blocking mitochondrial pyruvate import in brown adipocytes induces energy wasting via lipid cycling. EMBO Rep 21, e49634, doi:10.15252/embr.201949634 (2020).

46 Lee, H. S. & Lee, J. Influences of Ketogenic Diet on Body Fat Percentage, Respiratory Exchange Rate, and Total Cholesterol in Athletes: A Systematic Review and Meta-Analysis. Int J Environ Res Public Health 18, doi:10.3390/ijerph18062912 (2021).

47 Sperry, J. et al. Glioblastoma Utilizes Fatty Acids and Ketone Bodies for Growth Allowing Progression during Ketogenic Diet Therapy. iScience 23, 101453, doi:10.1016/j.isci.2020.101453 (2020).

48 Lee, J., Choi, J., Scafidi, S. & Wolfgang, M. J. Hepatic Fatty Acid Oxidation Restrains Systemic Catabolism during Starvation. Cell Rep 16, 201–212, doi:10.1016/j.celrep.2016.05.062 (2016).

49 Popow, J. et al. FASTKD2 is an RNA-binding protein required for mitochondrial RNA processing and translation. RNA 21, 1873–1884, doi:10.1261/rna.052365.115 (2015).

50 Perks, K. L. et al. PTCD1 Is Required for 16S rRNA Maturation Complex Stability and Mitochondrial Ribosome Assembly. Cell Rep 23, 127–142, doi:10.1016/j.celrep.2018.03.033 (2018).

51 Fleck, D. et al. PTCD1 Is Required for Mitochondrial Oxidative-Phosphorylation: Possible Genetic Association with Alzheimer’s Disease. J Neurosci 39, 4636–4656, doi:10.1523/JNEUROSCI.0116-19.2019 (2019).

